# Quantitative Structural Analysis of Influenza Virus by Cryo-electron Tomography and Convolutional Neural Networks

**DOI:** 10.1101/2021.12.09.472010

**Authors:** Qiu Yu Huang, Kangkang Song, Chen Xu, Daniel N.A. Bolon, Jennifer P. Wang, Robert W. Finberg, Celia A. Schiffer, Mohan Somasundaran

## Abstract

Influenza viruses pose severe public health threats; they cause millions of infections and tens of thousands of deaths annually in the US. Influenza viruses are extensively pleomorphic, in both shape and size as well as organization of viral structural proteins. Analysis of influenza morphology and ultrastructure can help elucidate viral structure-function relationships as well as aid in therapeutics and vaccine development. While cryo-electron tomography (cryoET) can depict the 3D organization of pleomorphic influenza, the low signal-to-noise ratio inherent to cryoET and extensive viral heterogeneity have precluded detailed characterization of influenza viruses. In this report, we developed a cryoET processing pipeline leveraging convolutional neural networks (CNNs) to characterize the morphological architecture of the A/Puerto Rico/8/34 (H1N1) influenza strain. Our pipeline improved the throughput of cryoET analysis and accurately identified viral components within tomograms. Using this approach, we successfully characterized influenza viral morphology, glycoprotein density, and conduct subtomogram averaging of HA glycoproteins. Application of this processing pipeline can aid in the structural characterization of not only influenza viruses, but other pleomorphic viruses and infected cells.

**Graphical abstract:** 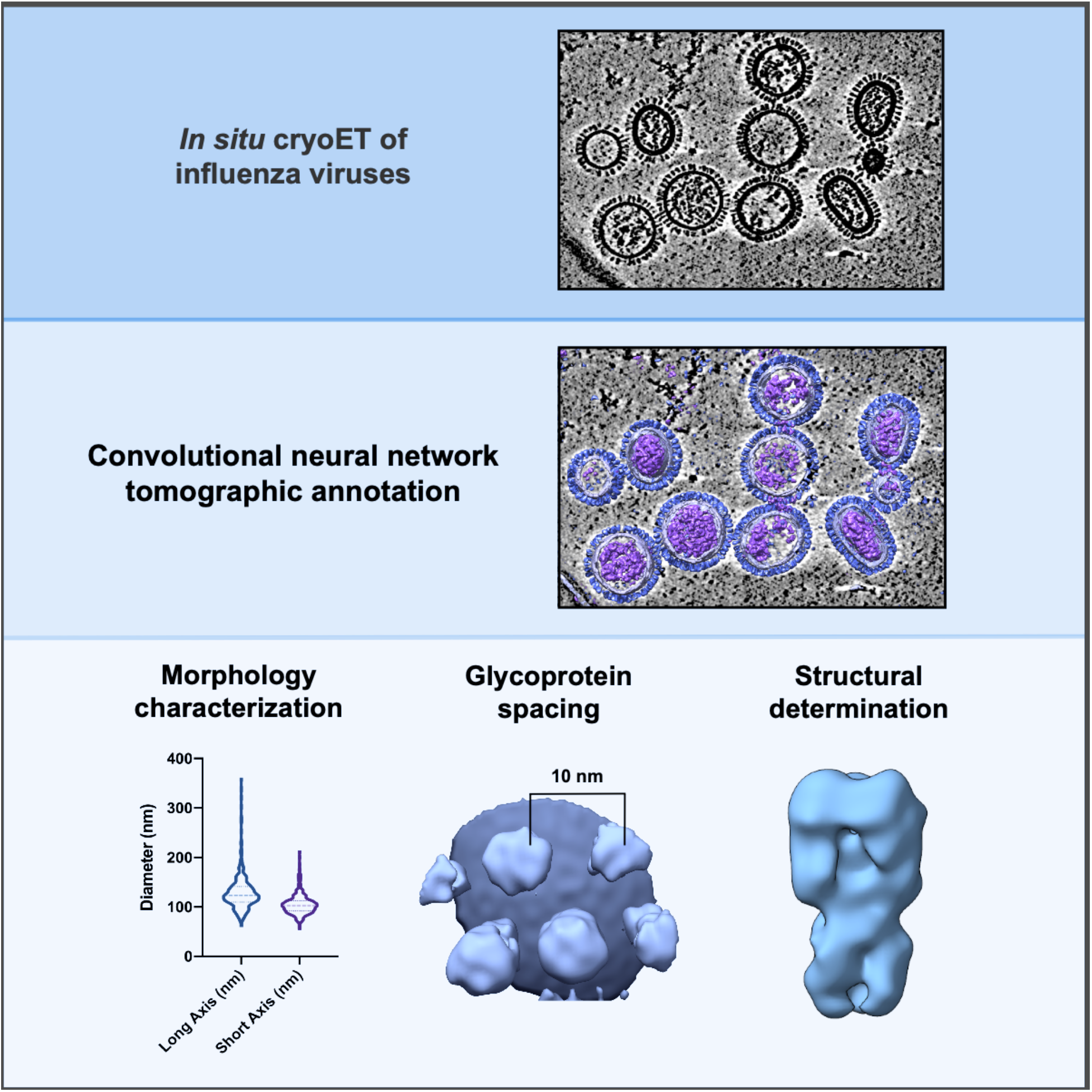

## Introduction

Influenza viruses are deadly human respiratory pathogens that pose severe public health threats. In the United States, the disease burden of influenza from the 2018-19 season was an estimated 35.5 million cases and 34,000 deaths (CDC, 2020; Krammer et al., 2018). While antiviral treatments and vaccine are available, influenza infections remain a public health concern due to genetic diversity and viral evolution. Influenza viruses are members of the *Orthomyxoviridae* family and are characterized by a segmented genome consisting of eight negative sense RNA strands (Krammer *et al*., 2018). This segmented genome mediates antigenic shift, which mediates cross-species transmission and enables animal viruses to infect humans. Moreover, influenza also evolves through antigenic drift, which is genetic diversity caused by RNA polymerase infidelity. Multifaceted influenza evolution pathways facilitate evasion from host immunity and necessitate the yearly reformulation of influenza vaccines (Corti and Lanzavecchia, 2013; Krammer *et al*., 2018; Laursen and Wilson, 2013). When there is an antigenic mismatch between vaccine and circulating strains, efficacy of protection can be low (Flannery, 2017).

Influenza viral particles express two essential and complementary membrane glycoproteins, hemagglutinin (HA) and neuraminidase (NA), on their surface. They perform essential roles in the viral infection cycle, and both glycoproteins recognize sialic acid on host cells (Gamblin and Skehel, 2010). HA mediates cellular entry by binding sialic acids on cell surface proteins and lipids to promote membrane fusion, while NA cleaves sialic acid from the cell surface to release newly budded viruses and spread to new cells (Gamblin and Skehel, 2010). HA is a homotrimer; each subunit contains a globular head domain or receptor-binding site, and a stem that contains the fusion peptide required for membrane fusion during cellular entry (Corti and Lanzavecchia, 2013). NA is a homotetramer that binds the host membrane via a thin stalk; each subunit contains an enzymatic active site that cleaves sialic acid for viral release (Gamblin and Skehel, 2010).

Influenza glycoproteins are asymmetrically distributed across the viral membrane, with an approximate HA/NA ratio of 5:1 (Harris et al., 2006; Vahey and Fletcher, 2019; Wasilewski et al., 2012). Due to their complementary roles, the balance of HA and NA is preserved despite constant evolution pressure. Their asymmetric distribution is hypothesized to promote efficient viral entry, budding, and motility; the HA/NA balance and is essential for preserving viral fitness during the emergence of antiviral resistance, host adaptation, and antigenic drift (Benton et al., 2015b; Benton et al., 2015a; de Vries et al., 2020; Gamblin and Skehel, 2010; Gaymard et al., 2016; Guo et al., 2018; Hooper and Bloom, 2013; Kosik et al., 2019; Mitnaul et al., 2000; Prachanronarong et al., 2019; Wagner et al., 2000; Xu et al., 2012). Further characterization of viral glycoproteins is important to tackle remaining fundamental questions on influenza organization and its role on viral fitness.

Influenza viruses are extensively pleomorphic. Virions from the same influenza strain that share identical genetic backbones can display a range of morphologies from small and spherical to elongated or filamentous (Dadonaite et al., 2016). While clinical isolates produce a mixture of spherical and filamentous particles that can reach up to 20 μm, extensively passaged lab isolates typically include small spherical virions of around 80-120 nm in diameter (Seladi-Schulman et al., 2013). Due to its extensive structure heterogeneity, 3D structural models of influenza viruses cannot be obtained using established methods such as X-ray crystallography nor single particle cryo-electron microscopy (cryoEM). Currently, characterization of influenza pleomorphism has been carried out using cryoET, a powerful cryoEM technique which has been used over the past two decades to characterize the 3D organization of pleomorphic influenza viruses *in situ* (Chen et al., 2019b; Fontana et al., 2012; Harris et al., 2006; Katz et al., 2014; Tran et al., 2016; Vijayakrishnan et al., 2013; Wasilewski et al., 2012). One main challenge that prevents detailed characterization of glycoprotein organization is the density of HA and NA glycoproteins on influenza virion surfaces. A spherical virus of 120 nm in width has around 300-400 glycoproteins (Harris *et al*., 2006), and a filamentous virus can express thousands of glycoproteins on a single virion (Wasilewski *et al*., 2012). CryoET studies have captured the intra- and inter-strain diversity of influenza viruses, but the intensive labor and manual input involved with data analyses have precluded large scale, quantitative analyses of pleomorphic viral morphology and glycoprotein organization.

To better quantify influenza morphology and glycoprotein distribution, we have incorporated convolutional neural networks (CNNs) in our cryoET workflow to annotate the main structural components of A/Puerto Rico/8/1934 (PR8), an egg-adapted H1N1 influenza A strain. As PR8 is a commonly used laboratory influenza strain and has been structurally characterized, we leveraged the known properties of this virus to develop cryoET processing pipeline for the structural characterization of influenza virions. Using this workflow, we have revealed further insights on influenza pleomorphism and surface glycoprotein organization.

## Results

### Influenza particles exhibit diverse morphologies

To evaluate the pleomorphic morphologies of influenza particles, 30 tomograms containing 311 PR8 virions were acquired and reconstructed (**Video S1, Video S2). Fig 1a**. shows a central slice through one tomogram. As expected, different morphologies can be detected in these tomograms with clear density (**Fig 1b**); Enveloped virions contain either solenoidal viral ribonucleoprotein (vRNP) complexes, (**Fig 1b** *I*), 7+1 vRNP complexes (**Fig 1b** *II*), or disorganized vRNPs (yellow arrow, **Fig 1b** *III*). The majority of the virions have a discernible M1 protein layer beneath their membrane, whereas a smaller subset does not contain the M1 density layer and only contain a lipid bilayer (**Fig 1b** *I*). On the surface of influenza viral particles, there are clear closely packed glycoprotein arrays. The sizes and shapes of the virions also varied greatly within the sample; from small and spherical (**Fig 1b** *I, II*), to oval (**Fig 1b** *V*), to elongated (**Fig 1b** *IV*), and irregularly shaped (**Fig 1b** *VI*).

**Fig 1.**
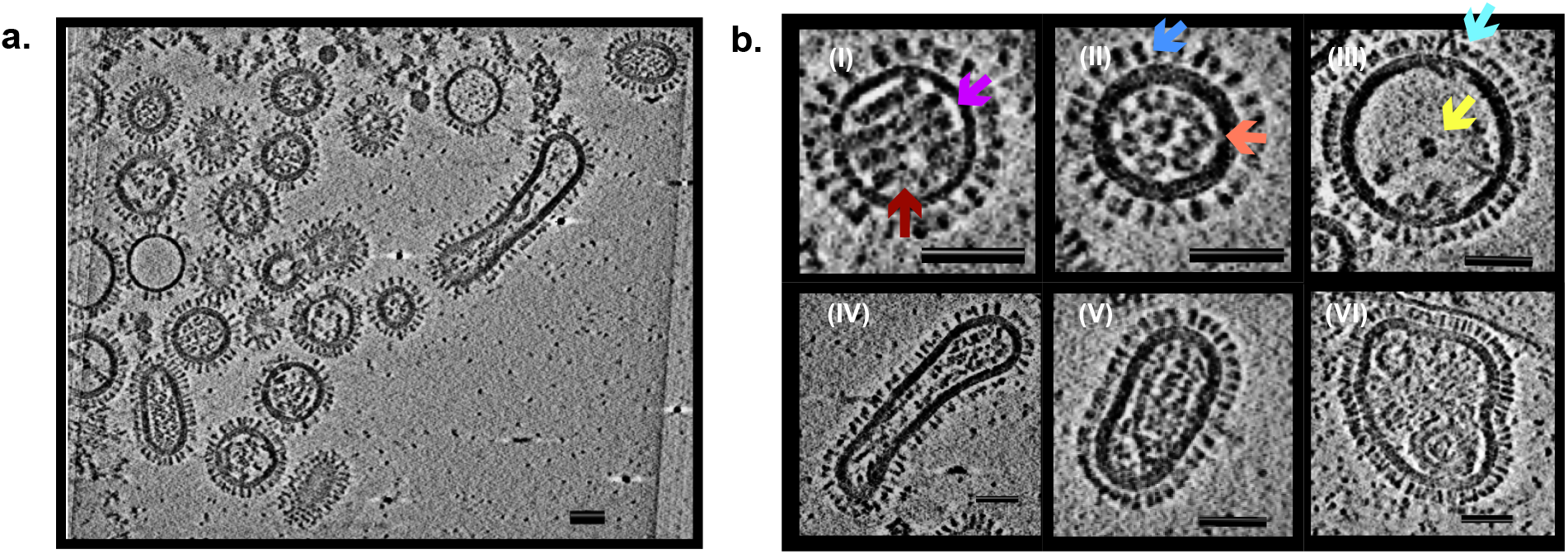
CryoET reveals diversity of IAV structures. **a. Central slice of representative tomogram. b. Examples of pleomorphic IAV particles** with (I) a solenoid vRNP complex organization (red arrow) and lipid bilayer without a M1 protein layer (magenta arrow); (II) lipid bilayers with M1 protein layer (orange arrow), HA (blue arrow), and an organized (7+1) vRNP core; (III) NA (aqua arrow), and a disorganized vRNP core (yellow arrow); (IV) elongated and (V) oval virions, and (VI) irregular virions. Scale bars = 50 nm.

### Convolutional neural networks (CNNs) accurately segment IAV structural components

To characterize the 3D morphology of influenza viruses at greater detail, the convolutional neural network (CNN) module implemented in the EMAN2 software (Chen et al., 2017) was used to train CNNs recognizing HA and NA glycoproteins, lipid bilayer, M1 protein complex, and vRNP complexes (**Fig 2, Fig S1)**. After individual networks were trained, each tomogram was segmented. The resulting segmentations were merged to create a multi-layer mask representing each structural component of influenza particles (**Fig 3**). To differentiate between virions where an M1 layer is present beneath the lipid bilayer (**Fig 3a**) and virions without the M1 protein assembly (**Fig 3b**), two networks were trained to recognize a thicker density layer and a thinner one, respectively. Despite the structural heterogeneity of pleomorphic influenza virions, training several CNNs corresponding to each virus structural component allowed for accurate and detailed visualization of 3D influenza virions (**Video S3, S4)**

**Fig 2.**
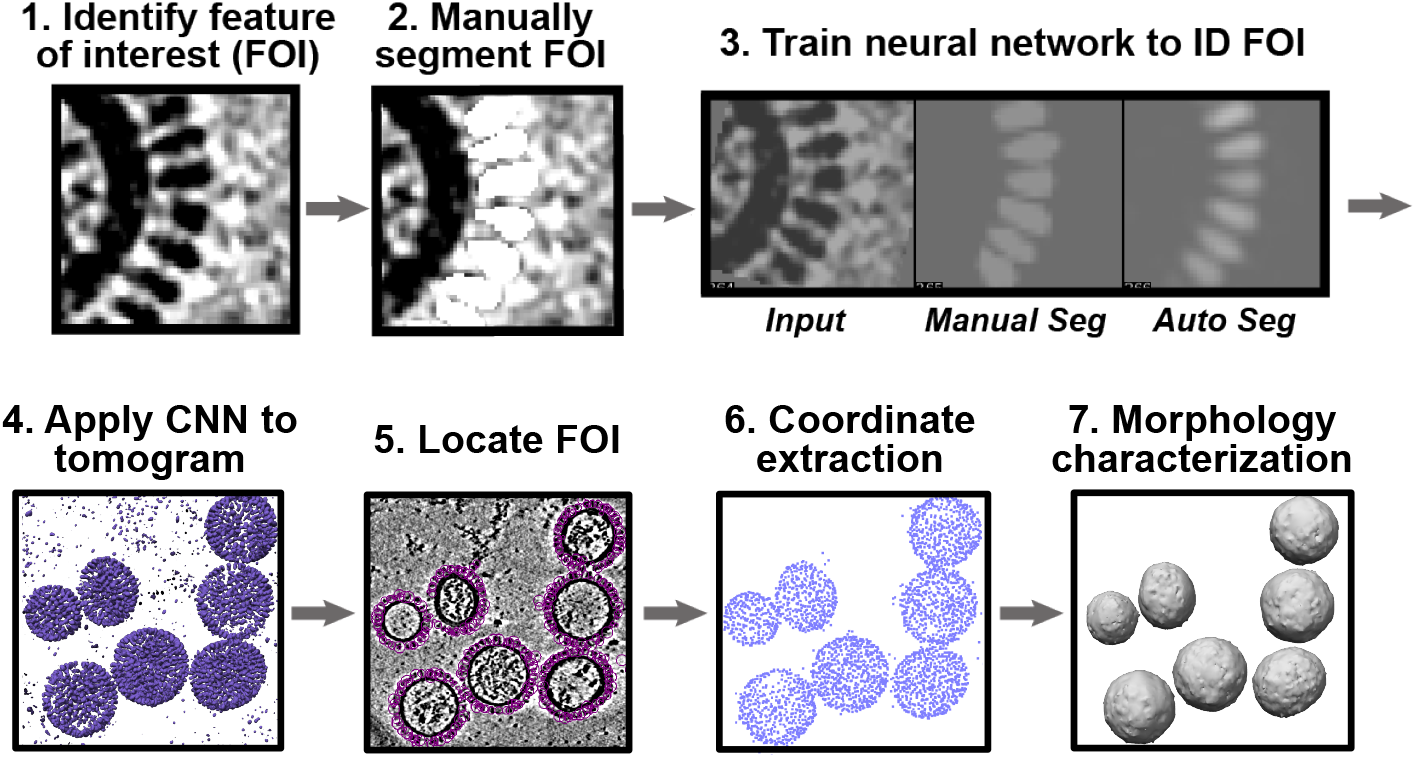
Example CNN-based analysis workflow of glycoproteins. Purple circles in (5) depict the locations of identified glycoproteins from a representative tomogram.

**Fig 3.**
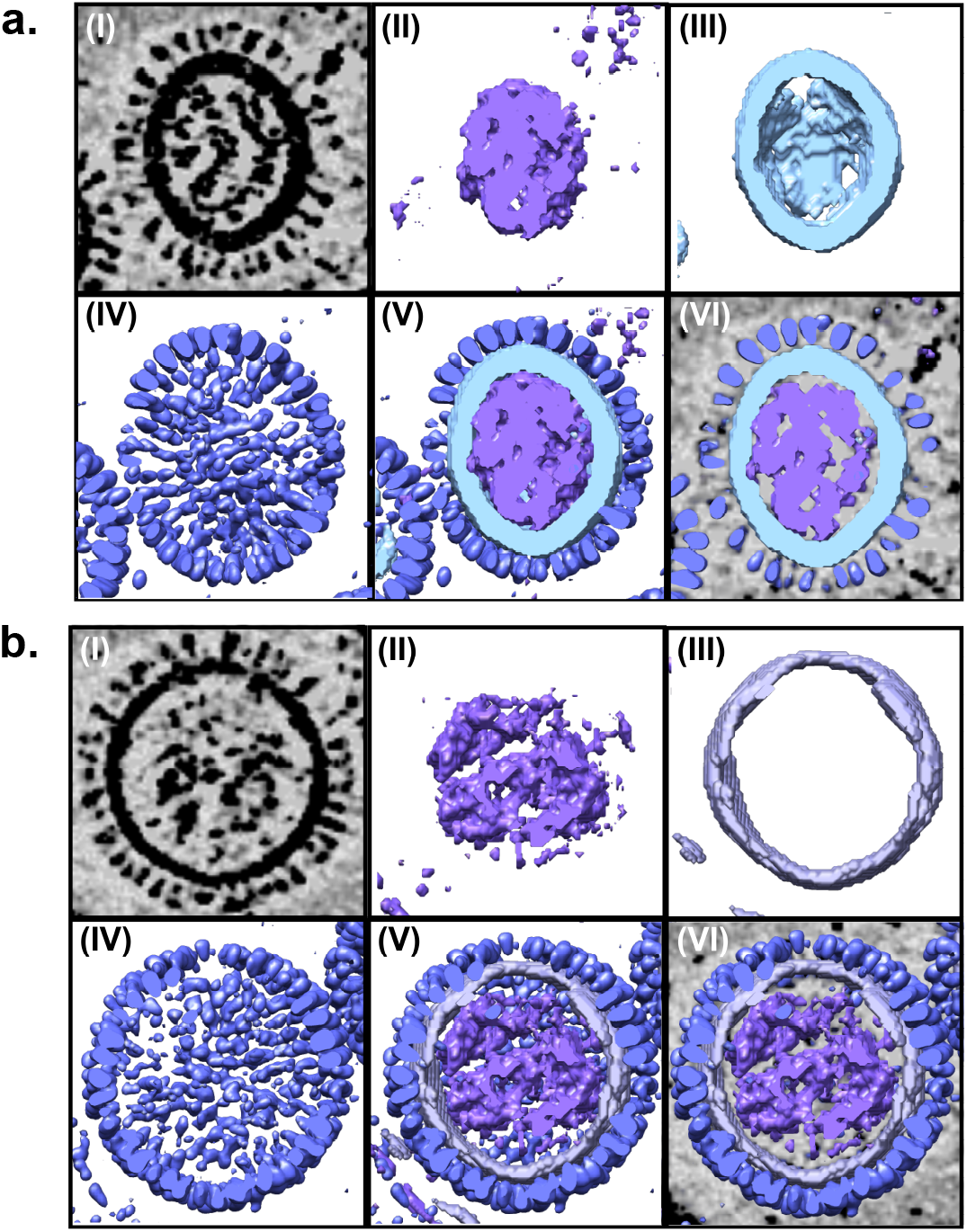
Convolutional neural network segmentation of influenza viruses. **a**. (I) Tomographic slice of a virus particle with vRNP complexes, M1 assembly, lipid bilayer, and glycoproteins. (II) Surface representation of vRNPs segmentation in purple. (III) Membrane segmentation in turquoise. (IV) Glycoprotein segmentation in blue. (V) Merged segmentation of the virus particle. (VI) Merged virus segmentation overlaid on tomogram. **c**. (I) Tomographic slice of a virus particle with vRNP complexes, lipid bilayer, and glycoproteins. (II) Surface representation of vRNPs segmentation in purple. (III) Membrane + M1 layer segmentation in lilac. (IV) Glycoprotein segmentation in blue. (V) Merged segmentation of the virus particle. (VI) Merged virus segmentation overlaid on tomogram. Surfaces of virus components were capped to visualize influenza internal structure beneath the glycoprotein and membrane layers.

Using CNN-guided segmentation, virions within tomograms can be accurately identified and isolated. The 3D structure of individual influenza particles can be visualized as triangulated surfaces. Importantly, this approach enables higher throughput morphological characterization of influenza particles, and properties such as presence or absence of the M1 protein, the size and shape of individual virions, and vRNPs presence and organization can be determined without extensive manual evaluations. Lastly, segmentation of individual virion components allows for flexible isolation of individual components, such as glycoproteins or vRNP complexes for further structural determination and interpretation.

To characterize the morphology of individual virions using this approach, the glycoprotein CNN was applied to each tomogram. As influenza viruses express a dense array of glycoproteins on virion surfaces, glycoprotein locations can be extrapolated as a measurement of virus size by connecting each point into a triangulated surface. The coordinates of glycoproteins identified by the CNN were extracted and modelled as point clouds (**Fig 2**). Each point cloud represents the 3D morphology of a single virion; thus, morphological analyses of influenza viruses were conducted based on the surface model each point cloud generated (**Fig 2**). This method provides an alternative approach from manually measuring viral axes in tomograms. Moreover, modelling each virion using glycoprotein coordinates examines 3D reconstructions instead of tomographic slices. This approach reduces potential biases concerning measuring virus size from tomographic slices due to the differential orientation of ice-embedded virions.

Additionally, the morphological profile of PR8 virions was characterized (**Fig 4**). In terms of size, influenza viruses were extensively pleomorphic. However, the vast majority of virions identified were spherical or oval in shape; the median axial ratio of this sample was 1.12 (**Fig 4a**). The long axes of PR8 virions ranged from 63 nm to 359 nm, with a mean of approximately 130 nm (**Fig 4a**). The short axis length was more narrowly distributed; they ranged between 56 and 211 nm with a mean of 105 nm (**Fig 4a**). Previous studies have shown that PR8 particles exhibited mostly spherical morphology, where only few particles are elongated or filamentous (Campbell et al., 2014; Chen *et al*., 2019b; Seladi-Schulman *et al*., 2013). These results recapitulate previous findings, which provides further confidence to this CNN-based pipeline. It was found that only twelve out of 311 particles have long axes more than twice the length of their short axis. The most prominent observation of a filamentous virion was of a particle that had a long axis which was 5.9 times longer than the short axis (**Table S1**).

**Fig 4.**
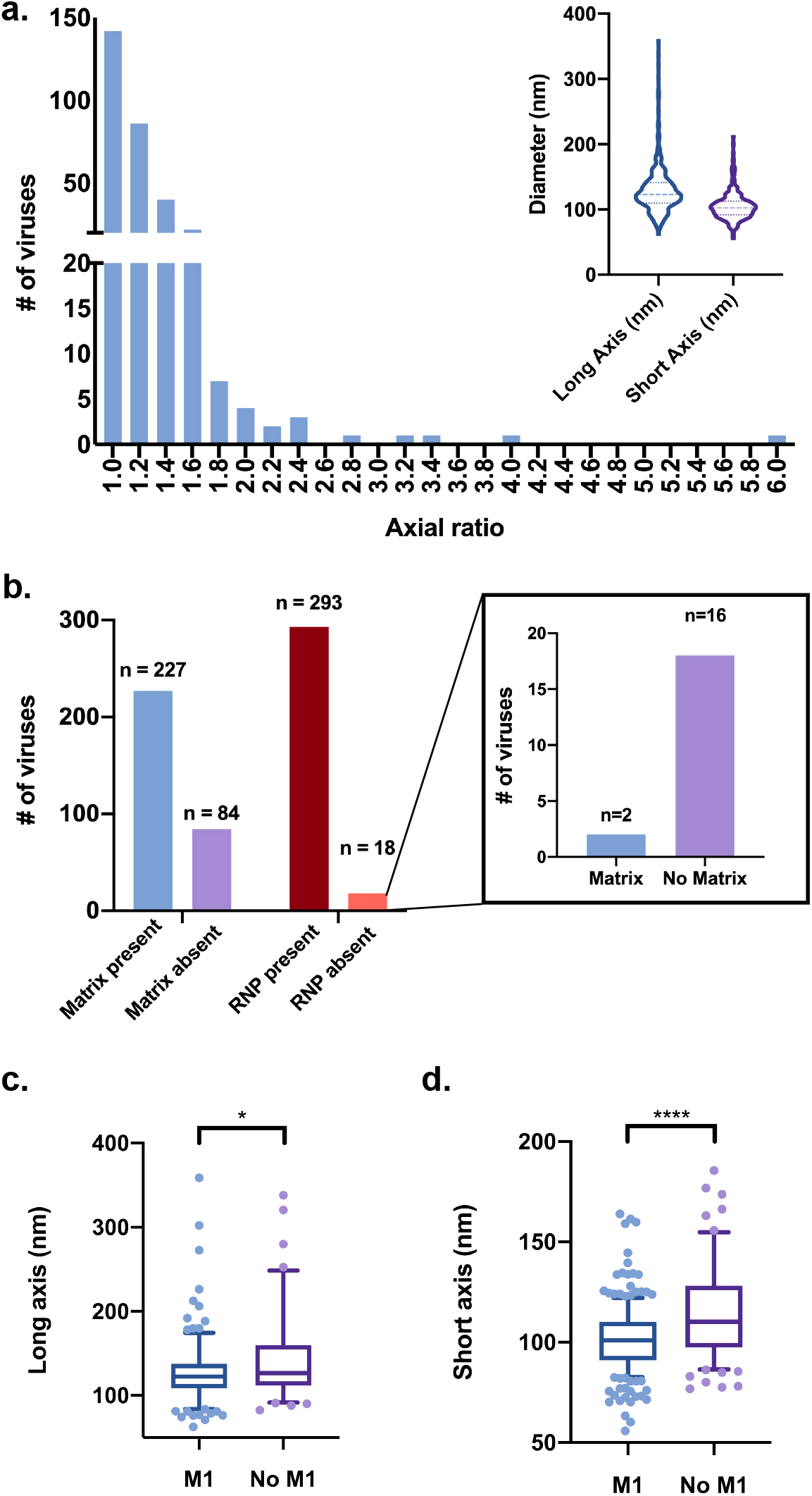
Morphological heterogeneity and M1 dependency of IAV particles. **a**. Histogram of axial ratios of IAV particles (n=311). **(Inset)** Violin plots for long and short axis lengths of virions. Middle dotted line indicates the median value; top and bottom dotted lines indicate interquartile range. Width of the plot correlates with the number of particles at that value. **b**. Presence of M1 assembly and vRNP in influenza virions. **c**. Box plot for long axis for virions with and without M1. Top and bottom lines indicate the interquartile range and the middle line median. Whiskers plot 5-95% of values and dots indicate each outlier. *P* = 0.0296 **d**. Box plot for short axis for virions with and without M1. Top and bottom lines indicate the interquartile range and the middle line median. Whiskers plot 5-95% of values and dots represent each outlier. *P* < 0.0001.

Additionally, the internal structural components of influenza particles were quantified (**Fig 4b**). Approximately a quarter (n=84) of virions did not have an intact M1 protein layer beneath the lipid bilayer (**Fig 4b**). M1 is an oligomeric protein important in viral assembly through interactions with both the membrane glycoproteins and the vRNP complexes (Ali et al., 2000; Peukes et al., 2020; Rossman and Lamb, 2011). It has been previous found to bind glycoproteins and vRNP complexes, as well as regulate filamentous particle formation of influenza viruses (Badham and Rossman, 2016; Elleman and Barclay, 2004; Rohlke and Stollman, 2012). Despite the larger size of filamentous influenza particles, several reports have shown filamentous virions only contain one copy of the genome (Badham and Rossman, 2016; Calder et al., 2010; Dadonaite et al., 2016; Noda et al., 2006; Vijayakrishnan et al., 2013). Within this sample, only a small portion of virions (n=18) lacked vRNP complexes (**Fig 4b**). The role of M1 in binding vRNP complexes for viral budding is further illustrated through our results, as 16/18 virions that lacked vRNPs also lacked M1, which underscores the importance of M1 binding on the incorporation of vRNPs into nascent virions. Furthermore, morphology quantification suggested that influenza particle size is modulated by M1. Size distribution of virions with the M1 protein is narrower in comparison to virions lacking M1 (**Fig 4e**). While the 25^th^ percentile and median long axis lengths for virions with and without M1 protein are comparable (**Table S1**), the distribution of viral axes length without the M1 protein is right-skewed. The 75^th^ percentile of M1-lacking viral long axis is 160 nm, whereas it is only 138 nm for viruses with M1 present. This pattern was more prominent for the short axis. PR8 short axis length shows an overall increase when the particle lacks a M1 protein layer; the median short axis of M1-absent particles is 10 nm greater than M1-containing virions (**Fig 4d**). Moreover, there were no virions with a short axis above 160 nm when the M1 protein was present, whereas the short axis of viruses lacking M1 extended up to 210 nm. These data further support that M1 helps mediate viral shape and morphology and underscores its role in regulating not only filament formation, but also the shape of non-filamentous influenza viruses.

### Influenza glycoproteins are evenly spaced on virion surfaces

Using CNN-guided particle picking, glycoprotein spacing on PR8 virions was quantified by extracting coordinates corresponding to HA and NA (**Video S5**). Coordinates from over 94,000 glycoproteins were exported, and distances between a single glycoprotein and its three closest neighbors were calculated. The average of these measurements was considered the inter-glycoprotein spacing for a glycoprotein. Despite the Consistent with previous reports where manual measurements found that each glycoprotein is located 10-11 nm from the next HA or NA (Harris *et al*., 2006; Wasilewski *et al*., 2012), the median inter-glycoprotein distance in this sample was measured as 9.6 ±3 nm (**Fig 5**). The 25^th^ percentile inter-glycoprotein distance was 8.6 nm and the 75^th^ percentile distance was 11.1 nm, suggesting that glycoprotein organization is tightly regulated on influenza virion surfaces. The appearance of outliers could indicate the presence of empty patches on influenza surfaces bereft of glycoproteins or partial virions captured within tomograms (**Fig S2**).

**Fig 5.**
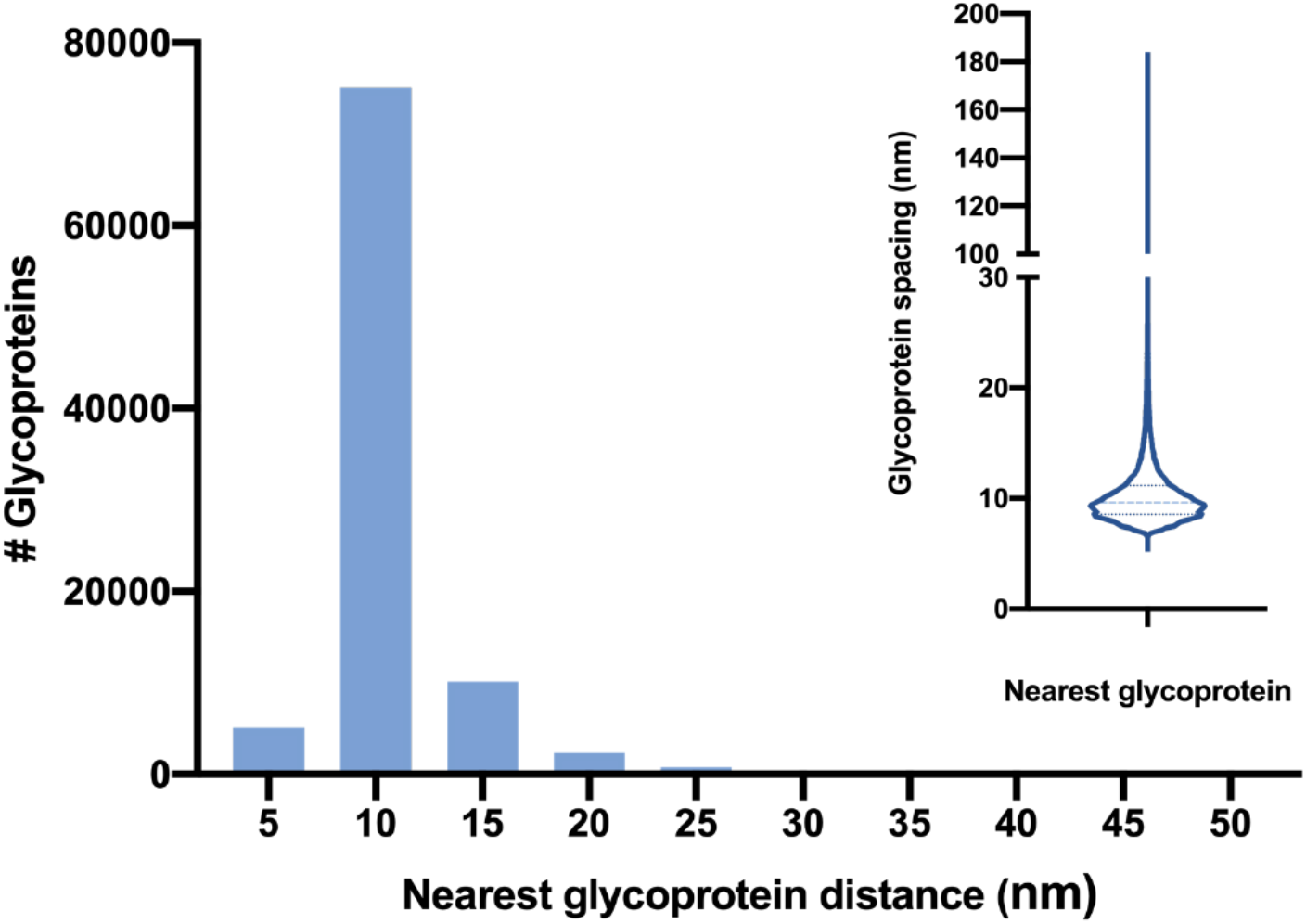
Glycoprotein spacing of IAV particles. **a**. Histogram of the average glycoprotein spacing between a viral glycoprotein with its three closest neighbours (n=94,281). **Inset**. Violin plot of the distribution of inter-glycoprotein spacing. Middle dotted line indicates the median spacing value; top and bottom dotted lines indicate interquartile range. Width of the plot correlates with the number of glycoproteins at that value.

The subtomogram average of a glycoprotein array confirmed the inter-glycoprotein spacing calculations. Subtomograms containing an array of glycoproteins were extracted for alignment and averaging (**Fig 6**). A cylindrical mask was applied on the extracted subtomograms post-alignment, which resulted in a clear array of three by two glycoproteins perpendicular to membrane density (**Fig 6a**). The lack of density between the glycoprotein stem and the membrane is most likely due to the flexible linker region. The median inter-glycoprotein spacing for the glycoprotein array was 10.5 nm, and the interquartile range was between 10 nm to 10.7 nm (**Fig 6b**). As glycoprotein coordinates were extracted from hundreds of virions, these data demonstrate that glycoprotein spacing is tightly regulated and well preserved on influenza virion surfaces.

**Fig 6.**
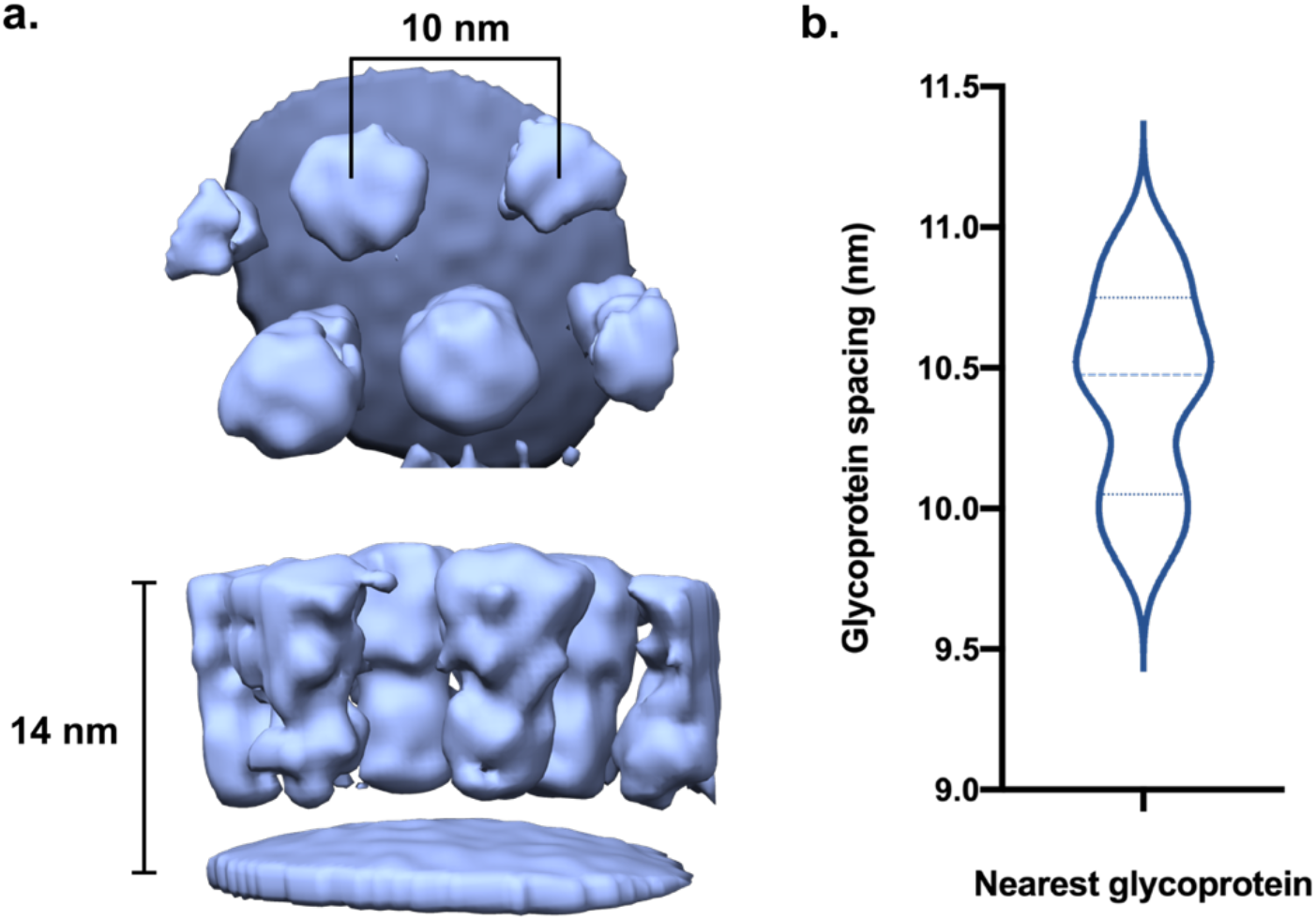
Glycoprotein spacing of aligned glycoprotein arrays. **a. (Top)** Top view of influenza glycoprotein array. **(Bottom)** Side view of glycoprotein array. **b**. Violin plot of the distribution of inter-glycoprotein spacing. Each value indicates the distance between a glycoprotein and its three closest neighbours. Middle dotted line indicates the median spacing value; top and bottom dotted lines indicate interquartile range. Width of the plot correlates with the number of glycoproteins at that value.

### Influenza glycoprotein spacing does not vary based on virion morphology

To investigate whether influenza particle morphology affects glycoprotein density, single virion glycoprotein spacing were calculated for spherical (1.0 < axial ratio < 1.2), oval (1.2 < axial ratio < 1.4), and elongated (axial ratio > 1.4) virions (**Fig 7**). Glycoprotein coordinates were extracted from individual virions. Between these populations, changes within glycoprotein spacing were small. The median inter-glycoprotein spacing, and interquartile range were comparable between all three morphologies (**Table S2**) While one-sample t tests between the populations reported a significant difference between the means for all groups, the differences in mean glycoprotein spacing vary by less than 0.5 nm on average and are likely too small to result in changes in viral infectivity and interaction with host. This finding further underscores the importance of regulating glycoproteins spacing on influenza viral surfaces.

**Fig 7.**
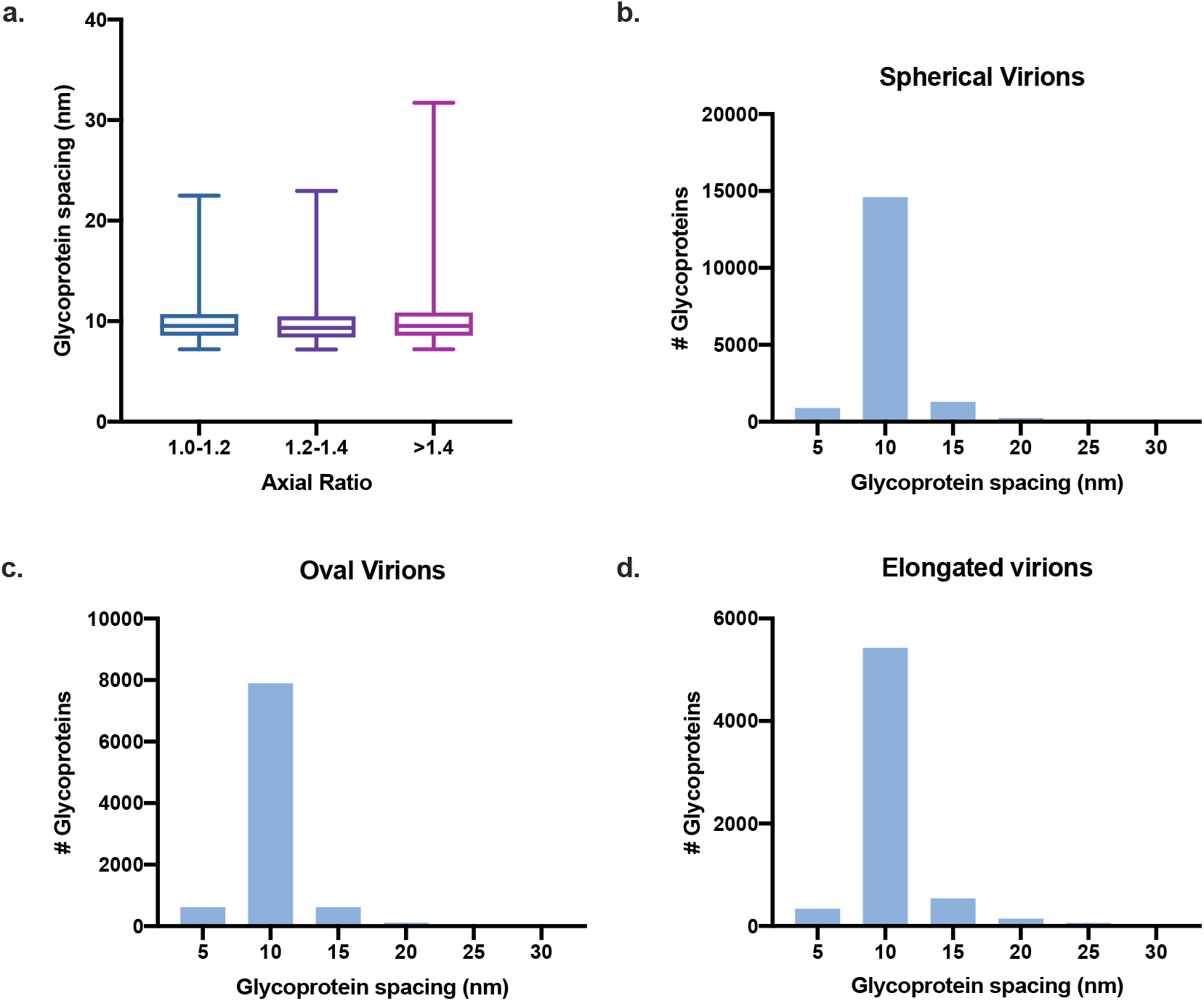
Glycoprotein spacing of IAV particles. **a**. Box plot comparing the inter-glycoprotein spacing distribution of spherical virions (n=17211), oval virions (n=9350), and elongated virions (n=6621). Top and bottom lines indicate the interquartile range and the middle line median. Whiskers indicate 1-99% of values. **b**. Histogram of inter-glycoprotein spacing on spherical particles between a viral glycoprotein with its three closest neighbours. **c**. Histogram of inter-glycoprotein spacing on oval particles between a viral glycoprotein with its three closest neighbours. **d**. Histogram of inter-glycoprotein spacing on elongated particles between a viral glycoprotein with its three closest neighbours.

### *In situ* influenza HA reconstruction

From the low-resolution glycoprotein array, *in situ* subtomogram averaging was conducted for focused refinement on one glycoprotein. As the extracted glycoprotein subtomograms were derived from a mixture of both HA particles and NA particles, reference-free subtomogram averaging was performed on glycoprotein particles without imposed symmetry for further classification analyses. Since initial alignments were performed with a large glycoprotein array, particle re-extraction was performed after five rounds of particle alignment to refine glycoprotein locations. This re-extraction eliminated the bottom 10% of particles according to the alignment score. After five further rounds of reference-free subtomogram averaging, three-fold symmetry emerged in the density map revealing a prefusion HA trimer (**Fig 8, Fig S3**). The resolved HA trimer was 13 nm in length and 8 nm in width. At an FSC=0.143, the resolution of the unmasked map was 13 Å, and map-to-model resolution was 17 Å (**Fig S4**). The map shows clear separation between globular head domains and stem domains of HA, as well as between the head domains of each HA monomer, and there was a central cavity between the three HA head domains (**Fig 8A**). Most likely due to linker flexibility, the membrane anchor portion of HA was not resolved in the map. Next, we compared the new *in situ* map with a previously determined crystal structure of H1 HA (Gamblin et al., 2004) (**Fig 8B**). Overall, the map agreed well with the structure, with a correlation score of 0.79. The separation between the head and stem domains of the HA crystal structure is accounted for by the central cavity between the globular head domains in the map.

**Fig 8.**
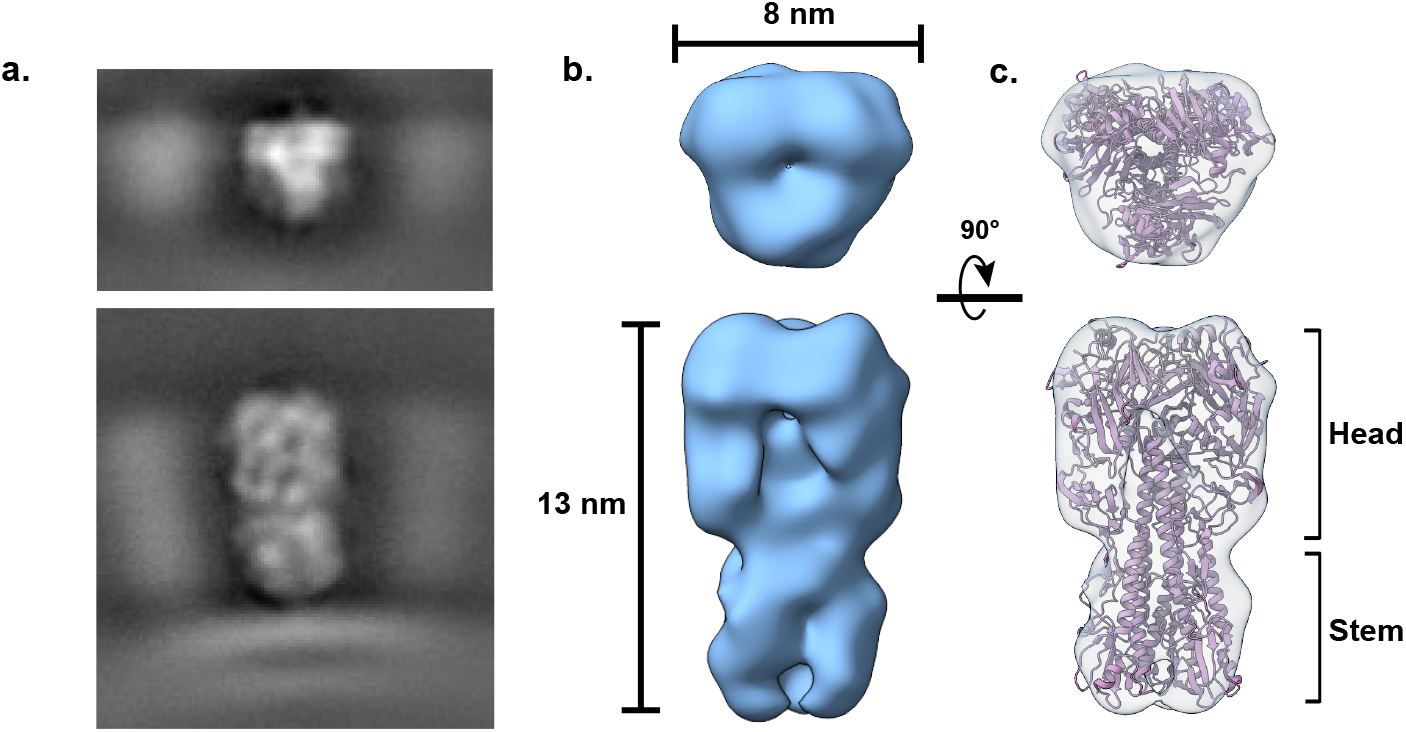
CryoET structure of influenza HA. **a**. (Upper) top projection image and (lower) side projection image of a HA trimer surrounded by neighbouring glycoproteins. **b**. (Upper) top and (lower) side view of a HA trimer. Density map was determined using reference-free subtomogram averaging from CNN-picked particles. **c**. (Upper) Top view and (lower) side view of a H1 crystal structure (PDB: **1RUZ**) docked into the HA subtomogram average using rigid body fitting.

## Discussion

Structural characterization of extensively pleomorphic influenza viruses has long been an arduous task. In this report, we developed a cryoET analysis pipeline incorporating CNN annotation for accurate morphological characterization of PR8 virions as well as *in situ* structural determination of a prefusion H1 HA trimer. This integrated methodology greatly improves processing throughput of tomographic data and locates individual influenza virions within tomograms.

Influenza viruses include multiple structural components including membrane-embedded surface glycoproteins HA and NA, a host-derived viral envelope, oligomeric M1 assemblies beneath the lipid bilayer, and vRNP complexes (Krammer *et al*., 2018). Our processing pipeline uses individual CNNs to annotate each of these components and dissects entire virions into distinct layers for further processing. As the CNN segmentation implemented in EMAN2 allows for combining several CNNs, this approach allows for flexible characterization of influenza viruses. Each particle is recognized as the composite of individual components instead of a virion of fixed size or shape. Therefore, influenza particles are readily identified despite morphological heterogeneity. Moreover, each viral structure is annotated independently, which allows for structural characterization of influenza virions without further subtomogram extraction and averaging (**Fig 3**). Size and morphological analysis can be fine-tuned using these annotations, including extraction of glycoprotein coordinates to calculate a 3D influenza virion model (**Fig 2**).

Our morphological analysis of PR8 particles showed that most virions were spherical or oval at a median axial ratio of 1.12. Moreover, the short axes of virions were tightly regulated, with the interquartile range of less than 20 nm. This finding was dependent on the presence of an M1 assembly; particles lacking M1 were on average more than 10 nm wider in the short axis. Moreover, both the long and short axes had wider distributions in the absence of M1. Our observations further support the role of M1 in regulating influenza morphology in non-filamentous strains.

While previous reports have characterized the glycoprotein density and spacing on influenza virion surfaces, the manual annotation required in locating glycoproteins have resulted in these calculations deriving from only thousands of glycoproteins from few virions. Here, harnessing our pipeline, we quantified the inter-glycoprotein spacing of close to 100,000 glycoproteins from hundreds of virions. We found that inter-glycoprotein spacing was the same regardless of particle morphology and the spacing was tightly regulated. Our pipeline can potentially detect altered glycoprotein expression in various strains in response to host immune pressures such as glycoprotein-targeting antibodies and antivirals. Likely, influenza surface organization is optimized for host receptor binding, entry, and egress. Maintenance of glycoprotein spacing may ensure the formation of proper clustering contacts with sialic acid on host cells (Ivanovic et al., 2013). The maintenance of glycoprotein spacing across several viral morphologies likely promotes influenza infection regardless of differences in shape or size. To verify the automated calculations, we performed subtomogram averaging on glycoprotein arrays extracted from the coordinates identified using CNNs (**Fig 6**). Our results corroborated well with each other and further demonstrates the ability of CNNs in extracting detailed structural data from pleomorphic viruses.

As the automated nature of particle picking enable enough glycoproteins for reference-free subtomogram averaging, we performed focused refinement on the glycoprotein array. To our surprise, a prefusion HA trimer emerged after reference-free subtomogram averaging. Using this approach, we were able to obtain a reconstruction of an *in situ* structure of influenza A HA.

In summary, we developed a tomographic analysis pipeline that enabled high throughput morphological analysis, quantification of glycoprotein density, and *in situ* structural determination of influenza viruses. This report demonstrates the ability of CNNs to aid in the characterizations of influenza A virus, a prototypical pleomorphic virus. Application of this pipeline towards other pleomorphic viruses such as SARS-CoV2 or HIV-1 may result in higher throughout processing of tomographic data and help interpret vast *in situ* structural information from purified viruses or infected cells.

## Author Contributions

Conceptualization, Q.Y.H, C.A.S, and M.S.; Methodology, Q.Y.H, C.A.S, and M.S.; Software, Q.Y.H; Formal Analysis, Q.Y.H; Investigation, Q.Y.H, K.S.; Resources, R.W.F, M.S., and C.X.; Writing – Original Draft, Q.Y.H, K.S., C.A.S, and M.S.; Writing – Review & Editing, Q.Y.H, K.S., J.P.W, C.A.S, and M.S; Funding Acquisition, D.N.A.B, J.P.W, R.W.F, C.A.S, and M.S.; Supervision, K.S., C.X., D.N.A.B, J.P.W, R.W.F, C.A.S, and M.S.

All authors approved of the final manuscript.

## Acknowledgments

The authors would like to acknowledge Drs. Nikolaus Grigorieff, Anne Jecrois, Florian Leidner, James Munro, and Nese Kurt Yilmaz for their advice and suggestions. The authors would also like to knowledge Drs. Muyuan Chen and Steven Ludtke for their helpful discussions surrounding the EMAN2 software package. This work was supported in part by Department of Defense W81XWH-14-PR 140464 and CAS was supported by R01 GM135919.

## Resource availability

### Lead contact

Further information and requests for resources and reagents should be directed to and will be fulfilled by the lead contact, Dr. Celia A. Schiffer.

### Materials availability

This study did not generate new unique reagents.

### Data and code availability

- The cryo-ET map of PR8 H1 is deposited at the EMDB and is publicly available as of the date of publication. Accession numbers are listed in the key resources table. Tilt-series and tomograms reported in the paper will be shared by the lead contact upon request
- Original Python scripts used for glycoprotein spacing calculations and virion surface model generation have been deposited on Github and is publicly available as of the date of publication. Github repository is linked in the key resources table.
- Any additional information required to reanalyze the data reported in this paper is available from the lead contact upon request.

## Materials and Methods

### Sample preparation

A/PR/8/34 (H1N1) influenza virus purified from embryonated chicken eggs was purchased from Charles River Laboratories and stored at -80 °C before use.

### Cryo-EM sample preparation and Tomographic data collection

3 μl of A/PR/8/34 (H1N1) influenza virus mixed with 1 μl of 5x BSA-coated 10 nm colloidal gold (Sigma-Aldrich) were applied to freshly washed and glow-discharged R2/2 holey carbon copper grids (Quantifoil). The grids were manually back blotted for 2 to 3s with filter paper, and rapidly frozen in liquid ethane using an EMS-2 rapid immersion freezer. The vitrified grids were stored in LN2. The specimen were imaged on a Thermo Fisher Scientific Titan Krios electron microscope operating at 300 kV with a K3 camera (Gatan, Pleasanton, USA) and a Gatan energy filter at a slit width of 20 eV. The tilt series were recorded using a dose-symmetric tilt scheme (Hagen et al, Implementation of a cryo-electron tomography tilt-scheme optimized for high resolution subtomogram averaging. 2017). The tilt series start at 0° in 2° or 3° increment at a magnification of 42,000 (corresponding to a calibrated 2.0873 Å of pixel size). And the tilt series range is limited to ±66°. Without a Volta phase plate, the defocus range is 3-4 µm. With a Volta phase plate, the defocus is 0.5 µm. The total dose used was less than 120 e/Å^2^. Data were acquired automatically under a low-dose mode using the SerialEM software (Mastronarde, 2005). The beam-induced motion of movie frames was corrected with IMOD software (Kremer et al., 1996).

### Tilt-series processing

Tilt-series alignment and tomogram reconstruction were performed with the EMAN2 software (Chen et al., 2019a). Alignment was conducted using an iterative 3D landmark approach, and Bin4 tomograms were reconstructed using a direct Fourier inversion algorithm for CNN training, annotation, and subtomogram averaging (Chen *et al*., 2019a). Final tomogram size was 2048×2048×512, at a sampling size of 8.35 Å/pixel. CTF estimation was carried out in EMAN2 for subsequent subtomogram averaging(Chen *et al*., 2019a).

### CNN-based tomogram annotation and segmentation

All CNN analysis was carried out using EMAN2’s semi-automated CNN module (Chen *et al*., 2017). For training, around 20-30 positive references and 150 negative references were selected for vRNP complexes, lipid bilayer + M1, and lipid bilayer alone. For robust recognition of influenza HA and NA, around 100 positive references and 200 negative references were used to train the glycoprotein CNN. To minimize false positives, a CNN was also trained to recognize carbon edges. After 100 rounds of training, individual networks were applied to each tomogram in the dataset and the annotations were merged to form a multi-layer mask. This process examines each voxel in the tomogram and assign the CNN annotation that has the highest value.

### Glycoprotein coordinate extraction

Coordinate extraction was carried out in two ways using CNN-based annotation. Particles were identified from the glycoprotein annotation mask using *e2spt_extractfromseg*.*py* or using the automated CNN particle selection tool *e2spt_boxer_convnet*.*py*. The coordinates of glycoproteins were exported and nearest neighbours analysis was performed using the Scikit-Learn library (Pedregosa et al., 2011).

### Morphology measurements

Initial virion morphology measurements were carried out in UCSF Chimera (Pettersen et al., 2004). Briefly, two measurements at the long and short axes of the particle were taken. After glycoprotein coordinates were extracted, coordinates from each tomogram were modelled as 3D point clouds using the Open3D library (Zhou et al., 2018). Outlier points were removed based on the standard deviation of a point’s distance from its closest neighbours from the average distances across the point clouds. Single virion point clouds were generated using the K-Means clustering algorithm implemented in the Scikit-Learn library (Pedregosa *et al*., 2011). Point clouds were exported as STL surfaces and each axis of the surface was automatically calculated in UCSF Chimera (Pettersen *et al*., 2004).

### Subtomogram averaging parameters

All subtomogram averaging steps were carried out in EMAN2(Chen *et al*., 2019a). To build an initial alignment model, 1000 extracted glycoprotein arrays were aligned and averaged; after five rounds of alignment, clear density of six glycoproteins perpendicular to a membrane emerged. The initial model was low pass filtered to 50 Å, and the full glycoprotein particle set containing 85,021 particles was aligned to this model with a cylindrical mask. The glycoprotein array subtomogram average after five further rounds of refinement was used for further refinement of glycoproteins. Based on the transformation matrix, glycoproteins were re-extracted. This step also discarded particles with the lowest 10% alignment scores as well as overlapping particles. At this stage, 76,519 particles were split into two sets based on the tomogram it was extracted, and independent refinement was carried out for both sets to ensure no overlapping particles were used for even/odd refinements. To reduce mask-related artefacts, no masks were applied at this stage. The subtomogram averages were carried out with the top 80% of particles according to alignment score. After five further rounds of 3D refinement, the Fourier Shell correlation was measured between unmasked maps from the two independent refinements. Rigid body fitting of a H1 trimer (PDB: 1RUZ) was performed in UCSF ChimeraX (Pettersen et al., 2021). Post-rigid body fitting, real space refinement was conducted using the *Phenix* software package; the refined structure was used to calculate a map-to-model FSC curve using the *Phenix* software package. (Adams et al., 2010).

### Quantification and Statistical Analysis

Statistical analyses were performed using GraphPad Prism 8.4.0. Data were represented as either violin plots or box and whisker plots. The number of virions and glycoproteins used in statistical analysis were indicated in the figure legends. Mann-Whitney U-tests were performed to compare the median axes lengths between particles with M1 and without M1. Welch’s T-test were conducted to compare mean glycoprotein spacing between virions of different morphologies.

### Visualization

UCSF Chimera (Pettersen *et al*., 2004) and UCSF ChimeraX (Pettersen *et al*., 2021) were used to visualize tomograms, tomographic segmentations, and subtomogram averages.

## Supplemental Figures

**Supplemental Figure 1.**
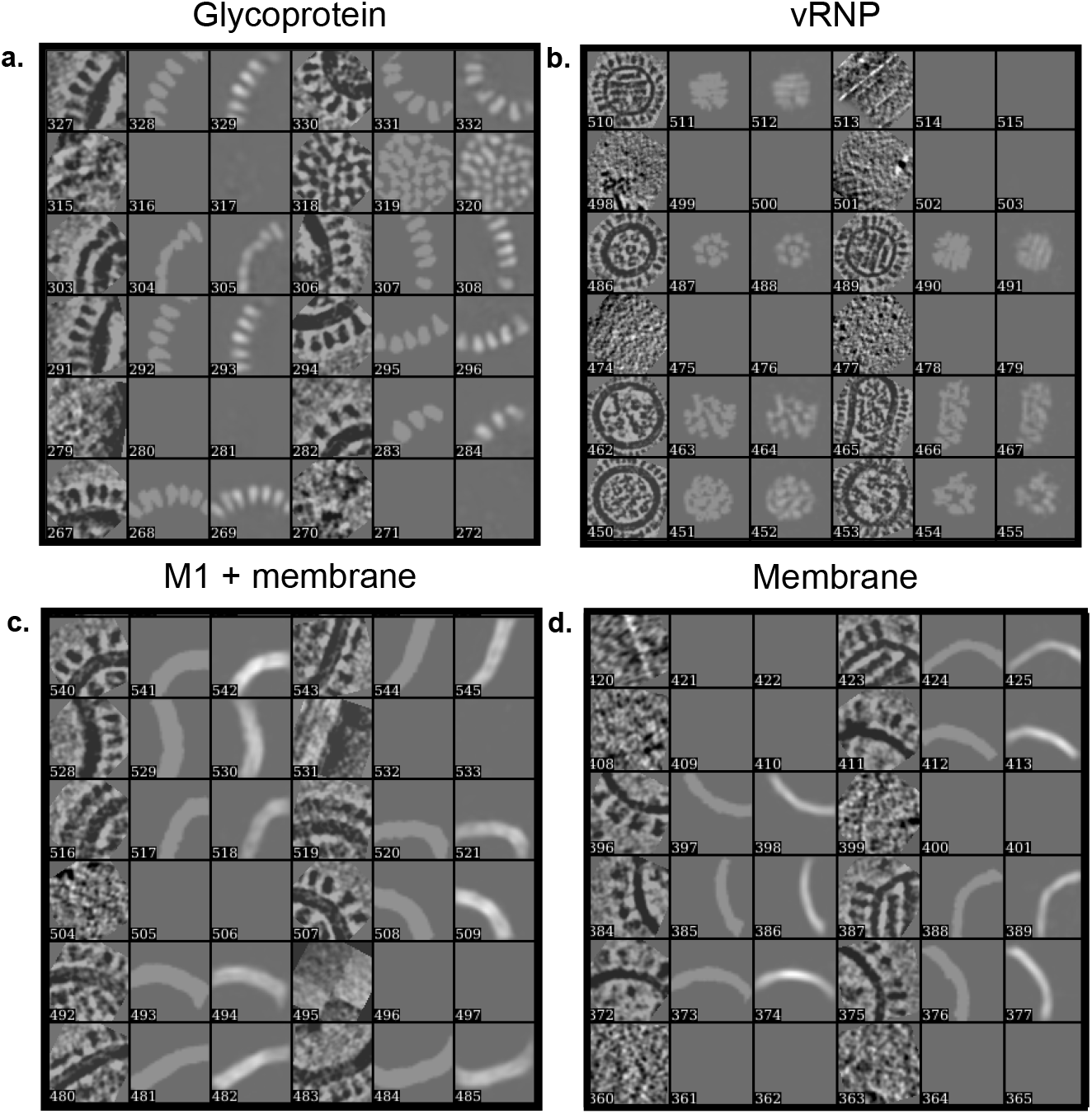
Training output of CNNs. Representative output from **(a)** glycoproteins **(b)** vRNP complexes **(c)** M1 + lipid bilayer **(d)** lipid bilayer alone are shown. The images represent the 2D tomographic slice, manual annotation of feature of interest, and automated annotation from CNNs. Samples without manual annotation represent negative samples. Related to Figure 2.

**Supplemental Figure 2.**
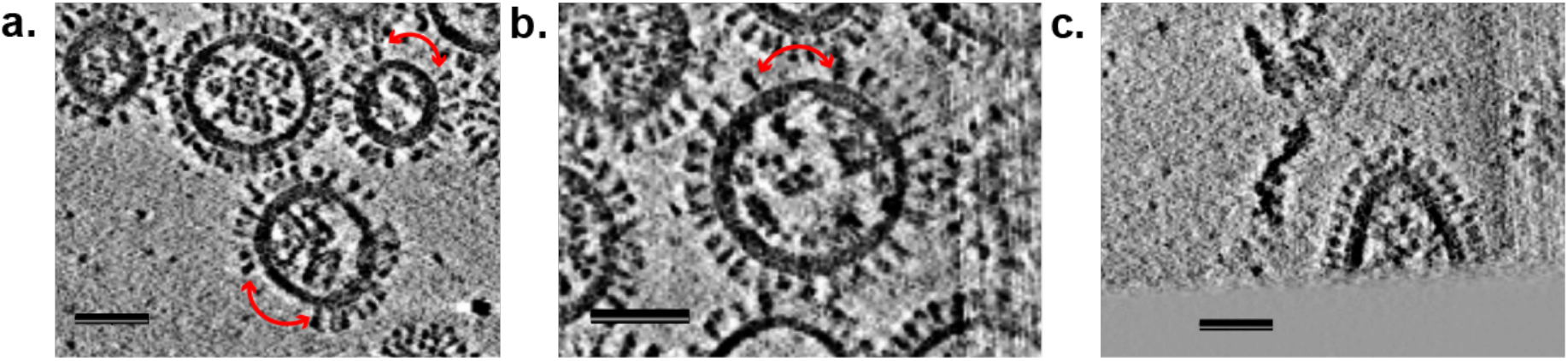
Influenza virions with empty membrane patches or cut off by grid edge. **(a)** and **(b)** show PR8 particles with membrane segments bereft of glycoproteins; sections are shown with red arrows. **(c)** is an example of a virion that was cut off. All scale bars are 50 nm. Related to Figure 5.

**Supplemental Figure 3.**
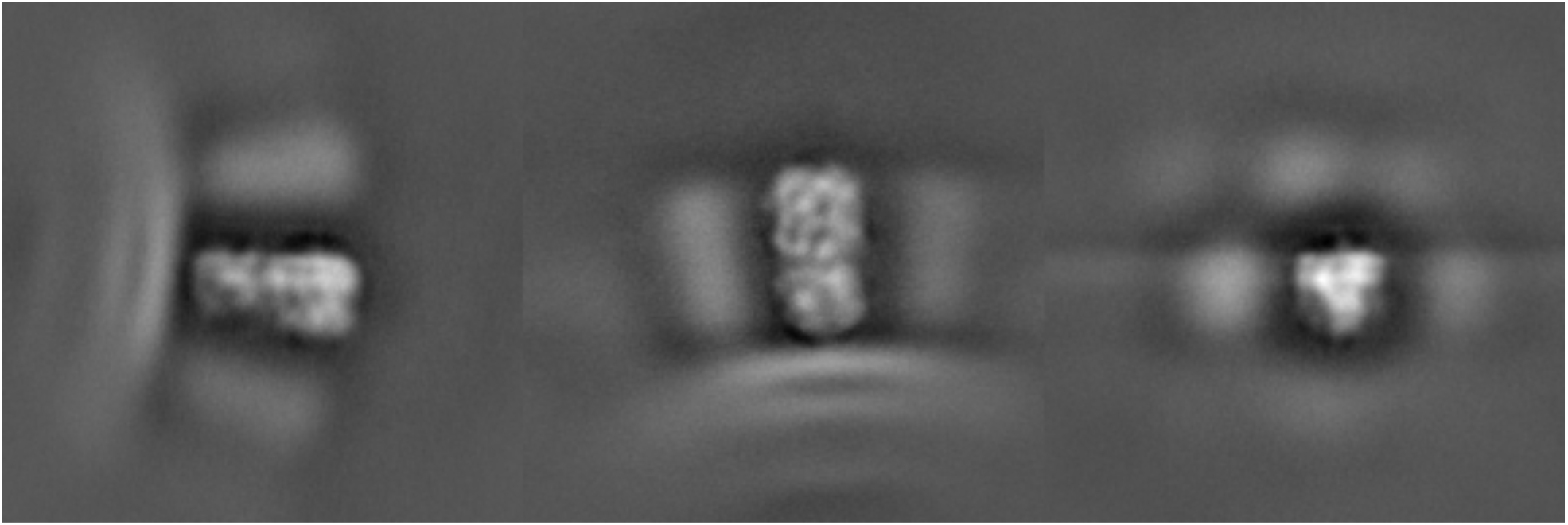
Images of unmasked HA trimer. X-, Y-, and Z-sections of the subtomogram average. Related to Figure 8.

**Supplemental Figure 4.**
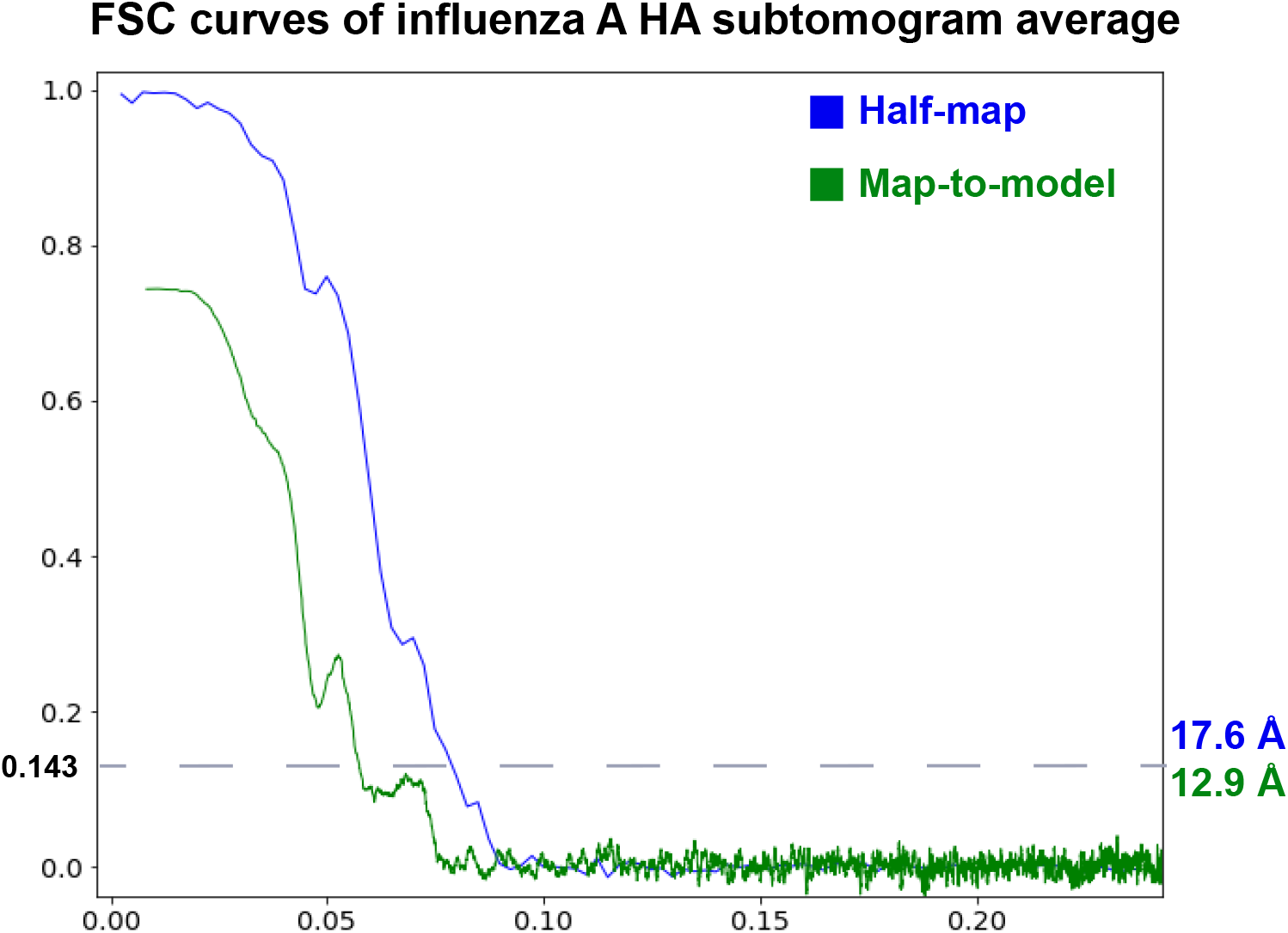
Fourier Shell Correlation plot for HA trimer. The resolution for the unmasked HA subtomogram average at FSC = 0.143 between the two individually refined sets is shown in blue. The map-to-model FSC between the subtomogram average and the fitted crystal structure (PDB: 1RUZ) is shown in green. Related to Figure 8.

**Supplemental Table 1.**
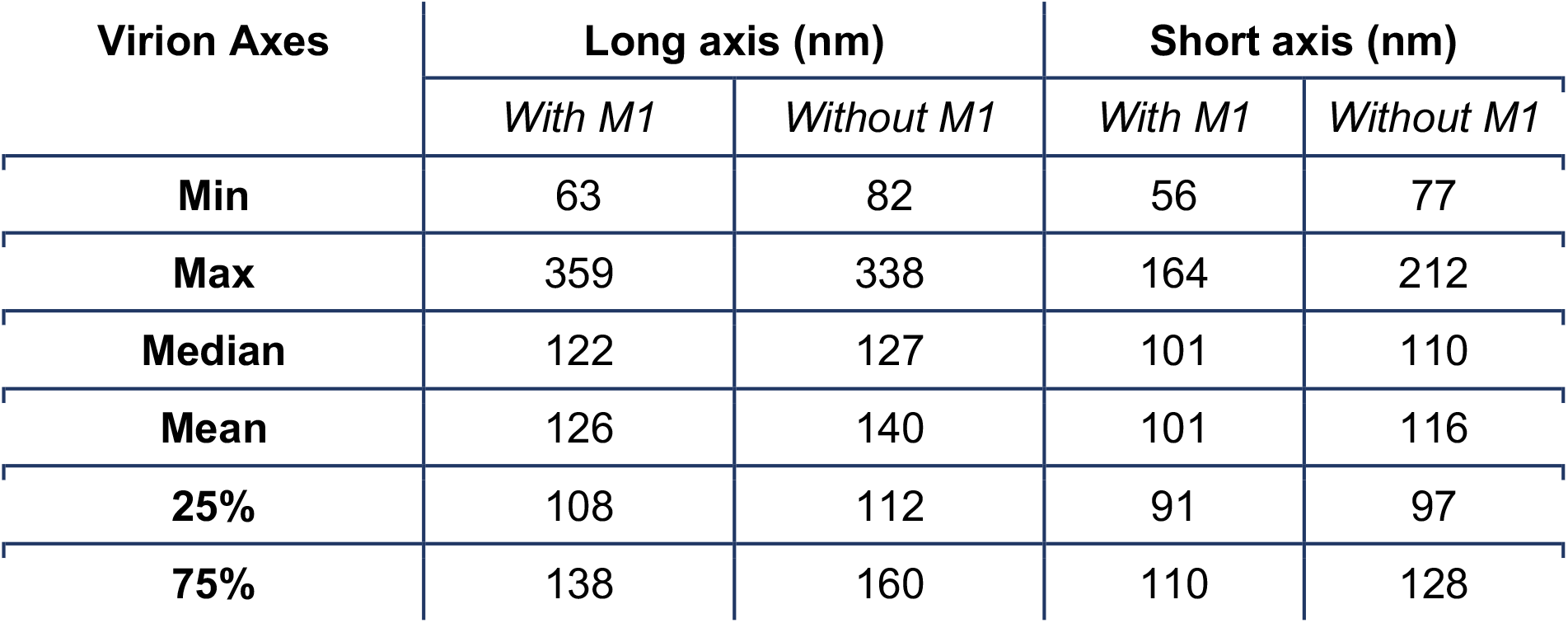
Virion axis measurements for influenza particles with M1 (228) and without M1 (84). Related to Figure 4.

| Virion Axes | Long axis (nm) |  | Short axis (nm) |  |
| --- | --- | --- | --- | --- |
|  | *With M1* | *Without M1* | *With M1* | *Without M1* |
| **Min** | 63 | 82 | 56 | 77 |
| **Max** | 359 | 338 | 164 | 212 |
| **Median** | 122 | 127 | 101 | 110 |
| **Mean** | 126 | 140 | 101 | 116 |
| **25%** | 108 | 112 | 91 | 97 |
| **75%** | 138 | 160 | 110 | 128 |

**Supplemental Table 2.** Inter-glycoprotein spacing measurements for PR8 virions of different morphologies. Measurements are in nm. Related to Figure 7.

| Glycoprotein spacing | Spherical (1.0-1.2) | Oval (1.2-1.4) | Elongated (> 1.4) |
| --- | --- | --- | --- |
| **Min** | 5.5 | 5.2 | 5.3 |
| **Max** | 87 | 93 | 99 |
| **Median** | 9.5 | 9.3 | 9.5 |
| **Mean** | 10 | 9.9 | 11 |
| **25%** | 8.6 | 8.4 | 8.6 |
| **75%** | 11 | 10 | 11 |

